# Inflammation in Areas of Fibrosis Precedes Loss of Kidney Function in Lupus Nephritis

**DOI:** 10.1101/2024.11.25.625225

**Authors:** Silvia Malvica, Paride Fenaroli, Chen-Yu Lee, Sarah Louis, Alessandra Ida Celia, Serena Bagnasco, Xiaoping Yang, Jeffrey B. Hodgin, Jill Buyon, Laurence Magder, Michelle Petri, the Accelerating Medicines Partnership: RA/SLE Network, Avi Rosenberg, Andrea Fava

**Author notes:** ***Corresponding author:*** Andrea Fava, MD, 1830 East Monument Street, Suite 7500 Baltimore, MD 21205, –. Drs Malvica, Fenaroli and Lee are co-first authors and contributed equally to this work. Drs Fava and Rosenberg are co-last authors and contributed equally to this work.

## Abstract

**Background:** Interstitial fibrosis in lupus nephritis (LN) is often infiltrated by immune cells but typically regarded as nonspecific “scar reaction.” This study aimed to investigate the relationship between inflammatory fibrosis and kidney disease progression in LN.

**Methods:** Interstitial fibrosis and tubular atrophy (IFTA) were scored in 124 LN kidney biopsies. Inflammation in areas of IFTA (i-IFTA) was graded 0-3 according to the Banff Classification of Allograft Pathology. Significant glomerular filtration rate (GFR) loss was defined as a decline of >15 ml/min at 3 years from biopsy. Immune cell phenotype was defined by serial immunohistochemistry (13-plex).

**Results:** IFTA was observed in 88/124 (71%) biopsies, and i-IFTA was identified in 76/88 (86%) cases. The distribution of i-IFTA grades was heterogenous across all IFTA grades. In patients with moderate-to-severe IFTA (>25%), the degree of i-IFTA was associated with a higher risk of significant GFR loss: 0/2 (0%), 1/3 (33%), 3/4 (75%), and 7/9 (78%) for i-IFTA grades 0, 1, 2, and 3, respectively (p = 0.028). Multiplexed histology revealed that i-IFTA was mostly composed of CD163+ macrophages and CD4 T cells, followed by CD8 T cells and granulocytes.

**Conclusion:** I-IFTA is frequently observed in LN and is dominated by macrophages and T cells. For patients with baseline IFTA >25%, the degree of i-IFTA emerged as a predictor of GFR loss. These data support the routine scoring of i-IFTA in LN due to its prognostic implications and nominate i-IFTA as a potential therapeutic target.

**LAY SUMMARY:** Scar tissue often contains immune cells, but we still do not fully understand their role. In lupus nephritis (LN), this is typically dismissed as “nonspecific inflammation”. However, our study analyzed kidney biopsies from 124 people with LN and found that inflammation in scarred areas may predict future kidney function loss. Specifically, we identified a type of immune cell, CD163+ macrophages, that may contribute to scarring and kidney damage. Our findings suggest that routinely assessing inflammation in scarred areas could help predict kidney health in LN patients and highlight a possible new target for therapies to prevent kidney damage.

## INTRODUCTION

Lupus nephritis (LN) is a major cause of morbidity and mortality in patients with systemic lupus erythematosus (SLE), leading to end-stage kidney disease (ESKD) in 10-30% of cases ^1, 2^. LN is currently classified into six classes based on the severity and distribution of glomerular histological features ^3^. Kidney histology also provides information on the degree of intrarenal LN activity (inflammation) and chronicity (damage) most commonly quantified by the NIH Activity Index (AI) and Chronicity Index (CI) ^3^. Current understanding of LN’s natural history suggests an evolution from an active, inflammatory phase to a resolving phase, where acute lesions may progress to fibrotic scarring through a process similar to wound healing. Within this framework, histological activity reflects potentially treatable lesions, while chronicity denotes irreversible damage. Chronicity in the diagnostic biopsy, but not activity, is the strongest predictor of future kidney function loss and progression to ESKD in LN ^4-9^. The CI is a composite score reflecting the extent of glomerulosclerosis, fibrous crescents, interstitial fibrosis, and tubular atrophy^3^. Interstitial fibrosis and tubular atrophy (IFTA) are the component of CI that most strongly predict negative renal outcomes in LN ^10-13^ as well as in other glomerular diseases such as IgA nephropathy ^14, 15^ and in the kidney allograft ^16^.

An inflammatory infiltrate in areas of IFTA (i-IFTA) is often observed but it is generally regarded as a non-specific feature in native kidney biopsies (scar reaction) rather than active disease requiring treatment. I-IFTA is also observed in non-immunological kidney disease such as diabetic nephropathy and hypertensive kidney disease, with similarly unclear clinical implications. In contrast, in kidney allograft pathology, i-IFTA is regarded as a lesion of immunological significance and is one of the histologic criteria of chronic active cellular-mediated rejection ^17^. However, the evidence supporting immunosuppressive treatment in this context remains inconclusive ^18, 19^

The clinical implications of i-IFTA in LN are poorly understood. The objective of this study was to investigate the association between inflammatory fibrosis and kidney disease progression in LN. Additionally, we characterized the immunophenotype of the inflammatory infiltrate in areas of IFTA.

## METHODS

### Study design and patient population

This study included patients enrolled in the Accelerating Medicines Partnership in RA/SLE cohort as previously described ^20-22^. In brief, SLE patients over 18 year of age were enrolled if they fulfilled the revised American College of Rheumatology (ACR) or the SLICC classification criteria for SLE, underwent a clinically indicated renal biopsy (defined as a UPCR >0.5 g/g)^20, 23-25^ and received a diagnosis of LN. Patient with a history of kidney transplant or pregnant at the time of biopsy were excluded. Only patients with available high resolution, whole digital images of the kidney biopsy light microscopy slides were included in study reported here. Clinical information, including serologies, were collected at the most recent visit before the biopsy.

Participants were treated for LN according to standard of care determined by their treating physician. Laboratory measurements were carried out in local laboratories with abnormal results defined as per the cutoffs of the laboratory.

Response status at week 52 was defined in patients with a baseline UPCR > 1 as follows: complete response (UPCR ≤ 0.5, normal serum creatinine or < 25% increase from baseline if abnormal, and prednisone ≤ 10 mg daily), partial response (UPCR > 0.5 but ≤ 50% of baseline value, normal serum creatinine or < 25% increase from baseline if abnormal, but prednisone dose could be up to 15 mg daily), or no response (UPCR > 50% of baseline value, new abnormal elevation of serum creatinine or ≥ 25% from baseline, or prednisone > 15 mg daily). The estimated glomerular filtration rate (eGFR) was calculated using the Chronic Kidney Disease Epidemiology Collaboration formula (CKD-EPI) ^26^. Significant loss of kidney function was defined as either reaching end-stage kidney disease (ESKD, eGFR <15 ml/min/1.73m^2^) or a decline of more than 15 ml/min/1.73m^2^ within 3 years (±3 months). This threshold is considered clinically relevant by guidelines and has been applied in similar studies of early CKD ^27^.

### Histological scoring

Renal biopsies were initially evaluated by a board-certified Pathologist at the institution where the biopsy took place and assigned histological classes, activity and chronicity indexes according to the International Society of Nephrology/Renal Pathology Society Classification. ^3, 28^. High resolution, whole digital slide images of formalin-fixed paraffin-embedded tissue stained with hematoxylin and eosin, periodic acid-Shiff, periodic acid silver methamine stain, and Masson trichrome stain were obtained using virtual microscopy (Concentriq, Proscia, Philadelphia, PA, USA; Aperio ImageScope v12.3.2.8013, Leica Biosystems, Wetzlar, Germany). The digital slides were centrally assessed for IFTA, and interstitial inflammation including i-IFTA by two operators and reviewed by a senior Renal Pathologist (SB). IFTA and i-IFTA were graded 0 to 3 based on the Banff Classification of Allograft Pathology (0: <5%, 1+: 6-25%, 2+: 26-50%, 3+: >50% and 0: <10%, 1+: 10-25%, 2+: 26-50%, 3+:>50%, respectively) ^17^.

### Statistical analysis

Descriptive statistics are presented as mean and standard deviation (SD) or median and interquartile range (IQR) for continuous variables, and frequencies for categorical variables. A two-tailed Student’s t-test or Wilcoxon’s test were used to compare continuous variables, and Fisher’s exact test was used to compare categorical variables where appropriate. Chi-squared for trend was used to associate inflammation in interstitial fibrosis and renal outcomes. Trends were tested by the Cochran-Armitage Trend Test and, in case or small sample size, confirmed by an exact test by permutation ^29^.

### Serial immunohistochemistry (sIHC)

Archival formalin-fixed paraffine-embedded (FFPE) were retrieved for serial immunohistochemical staining. In brief, sIHC involves multiple cycles of staining, destaining and restaining with high resolution images obtained with each cycle. Tissue sections (5 µm) were deparaffinized and subjected to antigen retrieval using citrate. Slides were blocked for peroxidases and incubated with primary antibodies, secondary HRP reagents, AEC-Red Chromogen, and Hematoxylin. Slides were decolored in 90% ethanol, stripped using antibody elution buffer, and sequentially cycled through the marker panel (CD3, CD20, CD138, PR3, CD4, CD8, CD14, CD16, CD163, CD66b, CD15, MPO, pankeratin). Images were captured at 40X magnification in each round.

Image files were imported into HALO-AI (version 3.5, Indica Labs) for deconvolution, registration, fusion and cell segmentation to create a registered composite image including all markers at single-cell level. Tissue regions including glomeruli and tubulointerstitium were initially identified by a DenseNet V2 AI classifier trained on a co-registered PAS and manually validated. Areas of IFTA were manually annotated (SM) and validated by an experienced renal pathologist (SB). Representative multichannel sIHC images were generated using pseudocolors.

Single-cell analysis was performed using R software (version 4.4.1, The R Foundation). Marker intensity was normalized using the centered-log ratio (CLR) transformation followed by scaling ^30^. Batch correction was performed using Harmony ^31^. The Seurat package (5.0.3) was used to perform dimensional reduction with PCA, followed by the construction of a KNN graph, and subsequent clustering using the Louvain algorithm.

Cell clusters exhibiting high pankeratin intensity were excluded to filter out renal epithelial cells. Clusters with immune marker levels exceeding 2 standard deviations above the mean were identified as candidate immune cell clusters for density analysis. The paired t-test was used to discover the significantly enriched immune cell types in IFTA or non-IFTA.

### Data attribution

The results published here are in whole or in part based on data obtained from the ARK Portal (arkportal.synapse.org).

### Ethical approval

In compliance with the Helsinki Declaration, all patients provided written informed consent approved by the respective institutional review boards and ethics committees of the participating sites for obtaining an extra renal core or leftover tissue and participation in follow-up visits. For this study, patients were included from Phase I and Phase 2 of AMP during which patients were followed longitudinally.

## RESULTS

### i-IFTA is a frequently observed feature of LN

We included 124 kidney biopsies from patients with LN who underwent a clinically indicated kidney biopsy and had a urine protein-to-creatinine ratio (UPCR) > 0.5 g/g. The clinical and demographic features are summarized in Table 1. Most patients were female (72.5%), with 47% identifying as Black, 23% as White, 13% as Asian, and 25% as Hispanic. At the time of biopsy, the majority had preserved kidney function and sub-nephrotic range proteinuria (mean eGFR 82.8 ml/min/1.73m2). Proliferative LN (pure class III, IV, or mixed with V) was present in 59% of patients, and NIH Activity Index (AI) and Chronicity Index (CI) scores varied widely (median AI 3 [range 3–18] and median CI 3 [range 0–9]).

**Table 1.** Clinicodemographic characteristics at the time of kidney biopsy.

|  |  | **i-IFTA score** |  |  |  |  |
| --- | --- | --- | --- | --- | --- | --- |
|  | **Overall** | **0** | **1** | **2** | **3** | **p** |
| **n** | 124 | 48 | 19 | 25 | 32 |  |
| **Age, years (mean (SD))** | 36.4 (12.7) | 36.9 (13.3) | 37.9 (15.3) | 32.40 (11.1) | 39.09 (11.8) | 0.442 |
| **Female (%)** | 79 (72.5) | 31 ( 70.5) | 11 ( 61.1) | 19 ( 79.2) | 18 ( 78.3) | 0.536 |
| **Race (%)** |  |  |  |  |  | 0.536 |
| **White** | 25 (22.9) | 10 ( 22.7) | 3 ( 16.7) | 7 ( 29.2) | 5 ( 21.7) |  |
| **Black** | 51 (46.8) | 19 ( 43.2) | 6 ( 33.3) | 12 ( 50.0) | 14 ( 60.9) |  |
| **Asian** | 14 (12.8) | 7 ( 15.9) | 4 ( 22.2) | 1 ( 4.2) | 2 ( 8.7) |  |
| **Other** | 19 (17.4) | 8 ( 18.2) | 5 ( 27.8) | 4 ( 16.7) | 2 ( 8.7) |  |
| **Hispanic (%)** | 27 (24.8) | 5 ( 11.4) | 7 ( 38.9) | 11 ( 45.8) | 4 ( 17.4) | 0.006 |
| **eGFR (mean (SD))** | 82.82 (28.82) | 93.35 (22.27) | 93.41 (30.49) | 78.74 (35.72) | 63.46 (22.74) | 0.005 |
| **UPCR (mean (SD))** | 2.07 (2.46) | 1.20 (0.95) | 1.46 (1.43) | 2.97 (3.32) | 2.91 (3.16) | 0.075 |
| **ISN class (%)** |  |  |  |  |  | 0.149 |
| **I/II** | 4 ( 3.6) | 4 ( 8.7) | 0 ( 0.0) | 0 ( 0.0) | 0 ( 0.0) |  |
| **Membr** | 42 (37.8) | 15 ( 32.6) | 8 ( 44.4) | 6 ( 25.0) | 13 ( 56.5) |  |
| **Mixed** | 29 (26.1) | 10 ( 21.7) | 5 ( 27.8) | 10 ( 41.7) | 4 ( 17.4) |  |
| **Prolif** | 36 (32.4) | 17 ( 37.0) | 5 ( 27.8) | 8 ( 33.3) | 6 ( 26.1) |  |
| **IFTA % (mean (SD))** | 16.50 (19.68) | 4.93 (11.40) | 15.57 (16.97) | 22.55 (19.63) | 28.31 (21.69) | <0.001 |
| **IFTA grade (%)** |  |  |  |  |  | <0.001 |
| **0** | 36 (29.0) | 36 ( 75.0) | 0 ( 0.0) | 0 ( 0.0) | 0 ( 0.0) |  |
| **1** | 58 (46.8) | 10 ( 20.8) | 14 ( 73.7) | 18 ( 72.0) | 16 ( 50.0) |  |
| **2** | 21 (16.9) | 1 ( 2.1) | 3 ( 15.8) | 5 ( 20.0) | 12 ( 37.5) |  |
| **3** | 9 ( 7.3) | 1 ( 2.1) | 2 ( 10.5) | 2 ( 8.0) | 4 ( 12.5) |  |
| **Activity Index (median [range])** | 3.00 [0-18] | 2 [0-18] | 3 [0-16] | 4 [0-15] | 3 [0-15] | 0.3 |
| **Chronicity Index (median [range])** | 3 [0-9] | 1 [0-6] | 3 [0-7] | 3.5 [0-9] | 5 [1-9] | <0.001 |

IFTA was observed in 88 of the 124 biopsies (71%), primarily mild: 58 patients (47%) had an IFTA score of 1, 21 patients (17%) had a score of 2, and 9 patients (7%) had a score of 3. I-IFTA was present in 76 of the 88 patients with IFTA (86%), and varying degrees of i-IFTA were distributed similarly across all IFTA score groups (**Figure 1**). Specifically, among the patients with IFTA, 12 (14%) had an i-IFTA score of 0 (<10% of IFTA), 19 (22%) had an i-IFTA score of 1 (10–25%), 25 (28%) had an i-IFTA score of 2 (25–50%), and 32 (36%) had an i-IFTA score of 3 (>50%).

**Figure 1.**
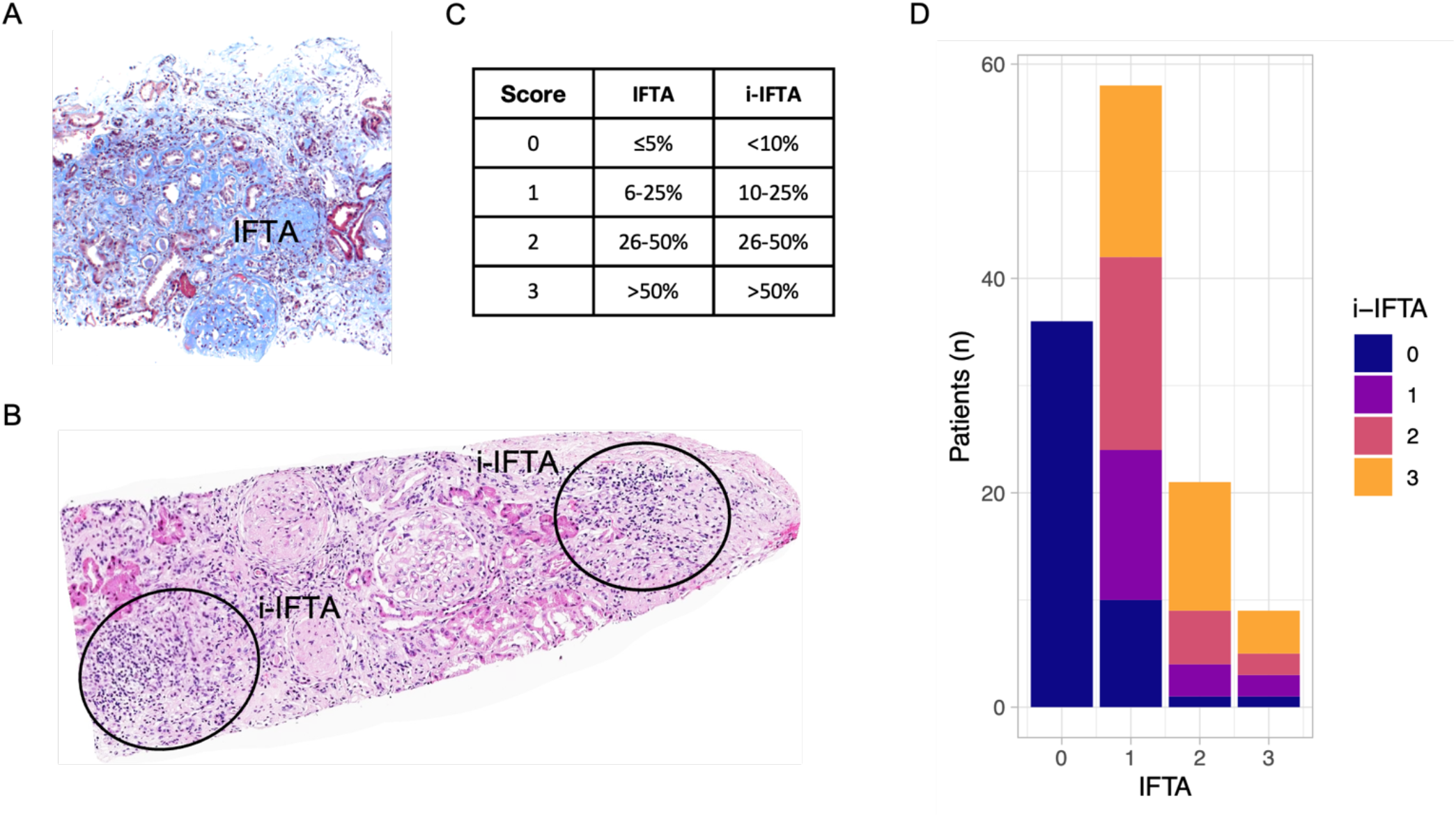
I-IFTA is frequently observed in LN. (A) Representative kidney biopsy showing extensive interstitial fibrosis (light blue stain by Trichrome) with interspersed tubular atrophy (IFTA). (B) Representative histological image showing an inflammatory infiltrated in areas of IFTA (i-IFTA). (C) Grading of IFTA and i-IFTA according to the percentage of fibrotic cortex for IFTA and the percentage of fibrotic cortex with inflammatory infiltrate for i-IFTA as per Banff criteria ^12^. (D) Distribution of i-IFTA grades (0-3) according to IFTA grades in 124 LN biopsies showing high frequency and heterogeneity in the degree of i-IFTA in patients with IFTA.

No significant associations were found between i-IFTA and age, sex, race, or treatment at time of biopsy (**Table 1**). Higher degrees of i-IFTA were associated with worse eGFR (p=0.005) and, numerically, with higher proteinuria (p=0.075). The distribution of i-IFTA was similar across most LN histological classes, with the exception of class II, where none of the four patients exhibited i-IFTA, despite one having an IFTA score of 3. There was no association between i-IFTA and the NIH Activity Index, which primarily focuses on glomerular inflammation. However, i-IFTA was significantly associated with higher NIH Chronicity Index scores (p<0.001).

### i-IFTA precedes loss of kidney function

Next, we tested whether i-IFTA was associated with future loss of kidney function. Longitudinal GFR data at 3 years from the index kidney biopsy were available for 69 patients (**Table 2**). Of these, 27/69 (39%) lost > 15 ml/min of GFR or developed ESKD.

**Table 2.** Clinicodemographic data and biopsy results of patients with the 3-year post biopsy follow-up according to GFR loss status.

|  | **GFR loss > 15ml/min or ESKD by 3 years** |  |  |
| --- | --- | --- | --- |
|  | No | Yes | p |
| **n** | 42 | 27 |  |
| **Age, years (mean (SD))** | 37.20 (14.32) | 37.40 (10.89) | 0.952 |
| **Female (%)** | 31 (73.8) | 22 (81.5) | 0.657 |
| **Race (%)** |  |  | 0.470 |
| **White** | 13 (31.0) | 7 (25.9) |  |
| **Black** | 18 (42.9) | 15 (55.6) |  |
| **Asian** | 3 ( 7.1) | 3 (11.1) |  |
| **Other** | 8 (19.0) | 2 ( 7.4) |  |
| **Hispanic (%)** | 12 (28.6) | 7 (25.9) | 1.000 |
| **eGFR (mean (SD))** | 87.58 (26.60) | 76.75 (30.49) | 0.176 |
| **UPCR (mean (SD))** | 1.87 (2.56) | 2.05 (1.73) | 0.781 |
| **ISN class (%)** |  |  | 0.757 |
| **I/II** | 2 ( 4.8) | 2 ( 7.4) |  |
| **Membr** | 17 (40.5) | 13 (48.1) |  |
| **Mixed** | 7 (16.7) | 5 (18.5) |  |
| **Prolif** | 16 (38.1) | 7 (25.9) |  |
| **Activity (median [range])** | 2 [0-18] | 2 [0-16] | 0.826 |
| **Chronicity (mean [range])** | 2 [0-7] | 5 [0-9] | 0.055 |
| **IFTA % (mean (SD))** | 13.7 (15.4) | 23.8 (25) | 0.056 |
| **IFTA grade (%)** |  |  | 0.1 |
| **0** | 13 (31.0) | 7 (25.9) |  |
| **1** | 22 (52.4) | 9 (33.3) |  |
| **2** | 5 (11.9) | 8 (29.6) |  |
| **3** | 2 ( 4.8) | 3 (11.1) |  |
| **i-IFTA grade** |  |  | 0.43 |
| **0** | 17 (40.5) | 9 (33.3) |  |
| **1** | 9 (21.4) | 3 (11.1) |  |
| **2** | 9 (21.4) | 7 (25.9) |  |
| **3** | 7 (16.7) | 8 (29.6) |  |

As expected, GFR loss was associated with the NIH Chronicity Index and higher IFTA (**Table 2**). Considering the relatively limited sample size, no other variable was significantly associated with loss of kidney function at 3 years. I-IFTA is graded according to the percentage of inflammatory infiltrates within areas of IFTA. Therefore, the total abundance of i-IFTA in the biopsy depends on the underlying amount of IFTA. For example, a mild degree of IFTA such as 10% (grade 1) could be infiltrated in 50% of its area (i-IFTA 3) but the total cortical area of i-IFTA would equal to 5%. In contrast, a severe degree of IFTA such as 50% (grade 3) could be infiltrated in 10% of its area (i-IFTA 1) thereby representing the same 5% of total cortical area of i-IFTA. To best analyze the clinical impact of i-IFTA and avoid this potential source of bias, we stratified patients according to the degree of IFTA into 2 groups: IFTA 0-1 and 2-3 (none to mild, and moderate to severe, respectively). The distribution of i-IFTA grades according to the degree of IFTA is illustrated in **Figure 2**. In patients with none to mild IFTA (grade 0-1, <=25%), the degree of i-IFTA was not associated with risk of GFR loss. In contrast, in patients with moderate to severe IFTA (grade 2-3, >25%), the degree of i-IFTA was associated with higher risk of eGFR loss (p for trend 0.0275). In patients with IFTA 2 or 3, absence of i-IFTA was associated with no increased risk of GFR loss. Conversely, the risk progressively increased with higher degrees of i-IFTA as 33%, 75%, and 78% for i-IFTA 1, 2, and 3, respectively.

**Figure 2.**
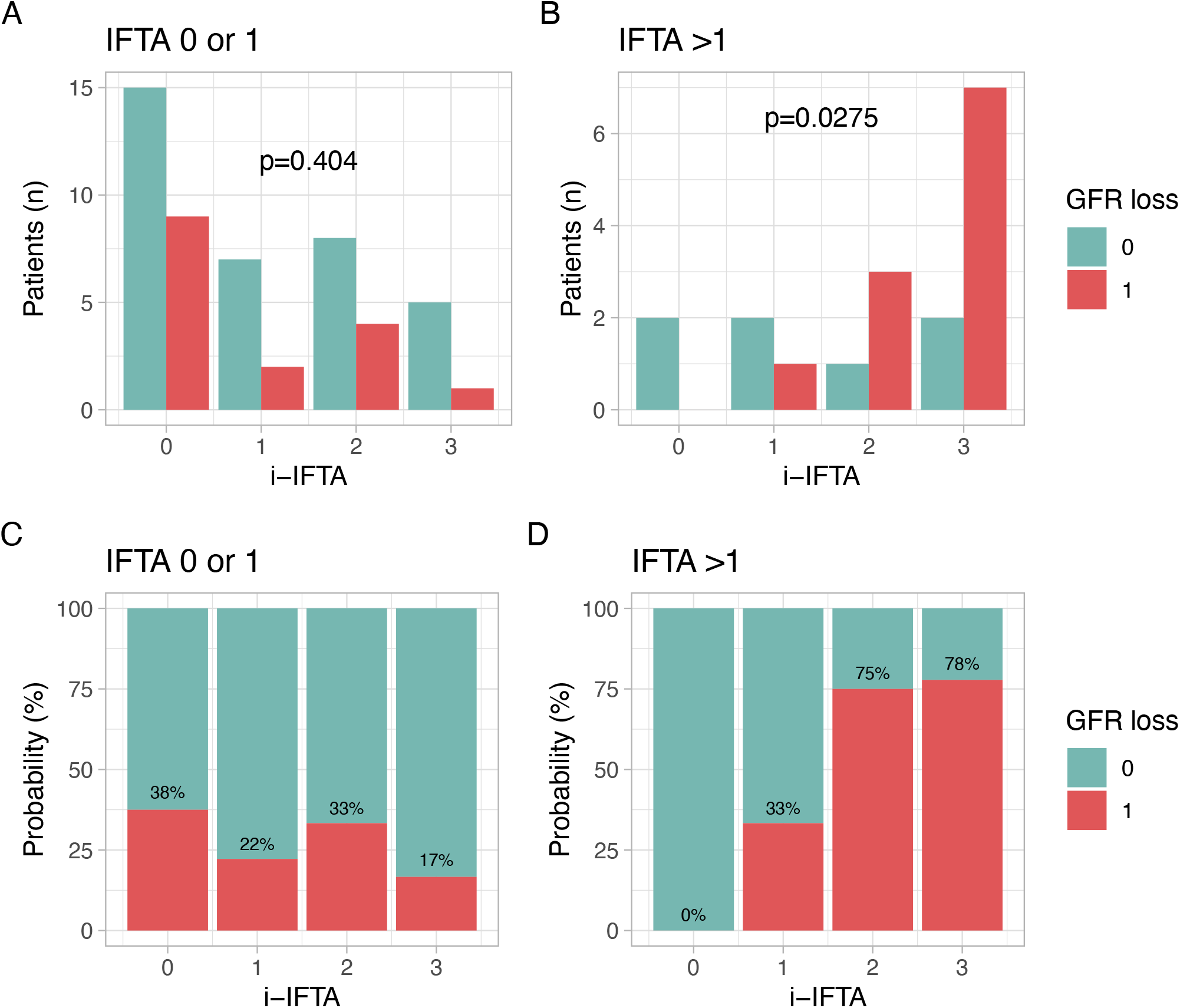
Risk of future GFR loss according to i-IFTA. Distribution of patients who developed significant loss of GFR (>15ml or ESKD) within 3 years of LN biopsy according to i-IFTA in patients with mild (A) or moderate-severe (B) IFTA. The results in A and B are reported as percentages in C and D.

### Phenotype of the immune infiltrate in i-IFTA

We employed serial immunohistochemistry to characterize the phenotype of the immune infiltrate in i-IFTA. We obtained 47,022 cells from 4 biopsies, including 8875 immune cells which clustered into 7 groups (**Figure 3A**). These included CD4+ T cells, CD8+ T cells, CD20+ B cells, CD163+ macrophages, and 2 clusters of granulocytes. Granulocytes were characterized by MPO expression with one cluster coexpressing PR3 and CD66b suggesting neutrophil lineage whereas the other showed variable PR3 expression but no CD66b suggesting a monocyte lineage (or atypical/nonpolylobate neutrophils based on morphology).

**Figure 3.**
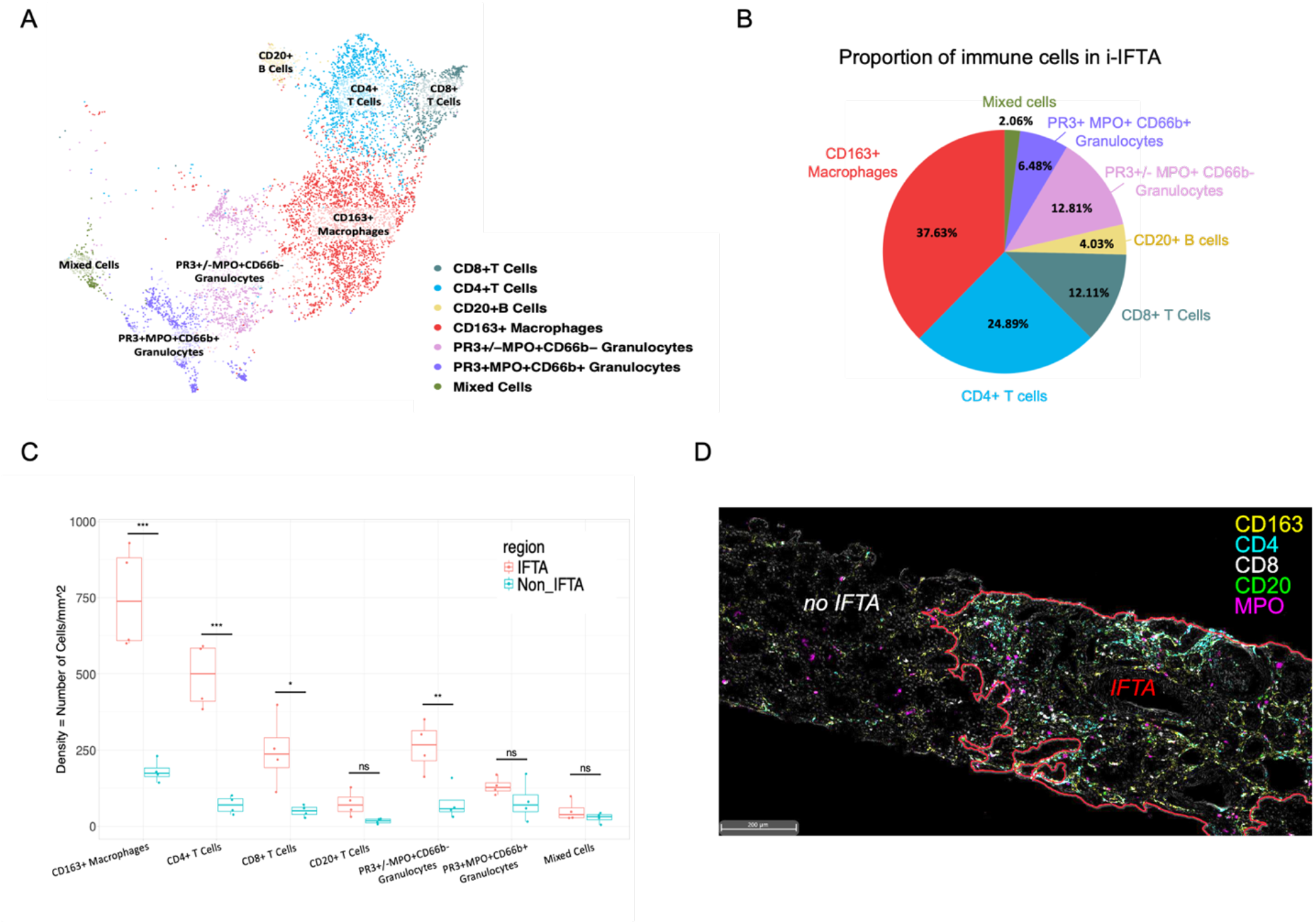
Immune cell composition of i-IFTA. Using 13-plex serial IHC, seven immune cell clusters were identified and annotated based on their marker expression. (A) UMAP of *** kidney-infiltrating immune cells (including all kidney regions) from 4 LN biopsies. (B) Pie chart showing the distribution of the immune cell infiltrate in i-IFTA. (C) Density of immune cell populations infiltrating areas of IFTA and tubulointerstitial regions without IFTA in 4 LN kidney biopsies. (D) Representative multichannel image showing the difference in immune infiltrate density in IFTA. Pseudocolors from 5 selected markers are shown. *Significance: *p = 0*.*05; **p < 0*.*05; ***p < 0*.*01; ns: not significant*.

The infiltrate in i-IFTA was heterogenous and included all cell types (**Figure 3B**). CD163+ macrophages (37.6%) and CD4 T cells (24.9%) represented the majority of the infiltrate in i-IFTA, followed by CD8 T cells, granulocytes, and B cells.

**Figure 3C** displays the density of each immune cluster in i-IFTA as compared to the infiltrate in tubulointerstitial regions without fibrosis. I-IFTA was characterized by a higher density of most immune cells, especially CD163+ macrophages and CD4+ T cells. A representative multichannel sIHC image illustrating the increase in CD163+ macrophages, CD4+ T cells, CD8+ T cells, and CD66b-granulocytes in IFTA is shown in **Figure 3D**.

## DISCUSSION

In this study, we demonstrated that i-IFTA is frequently observed in LN and showed substantial heterogeneity in its severity. I-IFTA is characterized by a heterogenous immune infiltrate, primarily composed of CD163+ macrophages and T lymphocytes. Although considered a nonspecific finding, the degree of i-IFTA emerged as a predictor of GFR loss, particularly for patients with baseline IFTA >25%.

IFTA is a well-established marker of injury severity and is a strong predictor of CKD progression in various kidney diseases, including in LN ^10-13^. Consistent with previous findings ^18^, our study confirmed that higher IFTA correlates with higher risk of GFR loss. The risk within IFTA was significantly influenced by the degree of the inflammatory infiltrate (i-IFTA). Higher i-IFTA predicted CKD progression in patients with moderate to severe IFTA, while the absence of i-IFTA despite IFTA was associated with no GFR loss. The limited number of patients in the last subgroup limits definitive conclusions. Nonetheless, these findings highlight the prognostic value of i-IFTA and support its routine inclusion and quantification in LN biopsy reports. The significance of i-IFTA in LN is further reinforced by previous studies showing that total tubulointerstitial inflammation, rather than inflammation in non-fibrotic regions, is linked to worse renal outcomes ^10, 32^. Together, these observations address a critical gap identified in the 2018 ISN/RPS revision to the histological classification of LN ^3^ and should inform the development of future scoring systems.

The association between i-IFTA and GFR loss reinforces the link between inflammation and fibrosis. This is further supported by the critical role of i-IFTA in the diagnosis of chronic active T-cell-mediated rejection ^17^ and association with poor kidney survival ^33-35^. Similar to other organs, fibrogenesis in the kidneys follows a common pattern where acute injury activates mesenchymal cells, fibroblasts, and pericytes, leading to a pro-inflammatory cascade ^36, 37^. This environment triggers myofibroblast proliferation, which promotes collagen secretion and extracellular matrix (ECM) deposition. Under normal conditions, this process is self-limited, with myofibroblasts undergoing apoptosis once the repair process is complete. However, if inflammation persists or injury is repeated, the repair process becomes unbalanced, resulting in excessive ECM deposition, parenchymal distortion, and functional decline ^38^. In the kidneys, this leads to scarring, nephron loss, and reduced renal function, with key contributors including myofibroblasts, macrophages, TGF-β signaling, epithelial to mesenchymal transition, and hypoxia ^39^. Given the association of i-IFTA with GFR loss, we hypothesize that i-IFTA contributes to fibrosis, possibly as a marker of an ongoing inflammatory process ^13, 40^ that could be targeted therapeutically to improve kidney outcomes. Animal models support this hypothesis, showing that the removal of macrophages after injury can prevent fibrosis ^41^. Previous repeat biopsy studies showed that kidney fibrosis in LN is partially reversible ^42^. Further studies should explore whether targeting i-IFTA can mitigate CKD progression in LN.

The immune infiltrate within i-IFTA was predominantly composed of CD163+ macrophages and CD4+ T cells, similar to the patterns observed in other fibrotic diseases and renal disorders such as IgA nephropathy and transplant rejection ^43-46^. In IgA nephropathy, CD68+ macrophage accumulation and CD3+/CD8+ T lymphocyte infiltration were linked to disease progression ^45, 47^. The association of CD163+ macrophages with decreased renal function is consistent across IgA nephropathy ^48^ and kidney transplantation ^49^, suggesting a shared mechanism of damage progression. CD163+ macrophages, which include alternatively activated (M2) pro-resolving and pro-fibrotic phenotypes, are thought to promote fibrogenesis by recruiting and activating myofibroblasts ^50^. Additionally, M2 macrophages may directly contribute to fibrosis through macrophage-myofibroblast transition^51^. Mechanistic studies demonstrate that M2 macrophages drive the transition from acute injury to chronic kidney disease and that their depletion reduces collagen deposition^50, 51^. The implication of macrophages the fibrosing process is observed across tissues such as in idiopathic pulmonary fibrosis ^24^, liver fibrosis ^52^ and cardiac neonatal lupus ^53^. In addition to macrophages, the presence of CD4+ and CD8+ T cells, and B cells along with granulocytes and monocytes implicates a broad interaction between the adaptative and innate immune system ^54^. Whether the response within IFTA is driven by a specific antigen as previously observed in interstitial inflammation^55^ remains to be determined.

We acknowledge several limitations of this study. The sample size restricted our ability to conduct subgroup analyses, such as stratifying by ISN class or assessing the impact of medication use. Furthermore, we could not determine the “induction” regimen for each patient. Important confounding factors such as hypertension control, medication adherence, or nephrotoxic events could not be fully accounted for. Additionally, key cell types involved in fibrosis, such as fibroblasts, dendritic cells, and plasma cells, were not quantified, limiting our ability to study cell-cell interactions systematically.

In conclusion, our findings establish i-IFTA as a negative prognostic marker in LN, predicting poor renal outcomes. The presence and severity of i-IFTA should be routinely reported in biopsy assessments due to its prognostic significance. Importantly, i-IFTA represents a potential therapeutic target, and future research should focus on whether interventions aimed at reducing interstitial inflammation can slow or prevent CKD progression in lupus nephritis and other fibrotic renal diseases.

## DISCLOSURE

The AMP is a public-private initiative as detailed under Acknowledgment. There were no conflict of interests related this work.

## DATA STATEMENT

The data generated by the AMP are publicly available at https://arkportal.synapse.org/.

## Acknowledgements

We would like to thank Jennifer Lee and Daksh Saksena for their support in the draft and review of the manuscript. This work was funded by the Plank Family Foundation and the Jerome L. Greene Foundation. The Hopkins Lupus Cohort is supported by R01-DK-134625. This work was supported by the Accelerating Medicines Partnership® Rheumatoid Arthritis and Systemic Lupus Erythematosus (AMP® RA/SLE) Network (AMP) in Rheumatoid Arthritis and Lupus Network. AMP is a public-private partnership (AbbVie Inc., Arthritis Foundation, Bristol-Myers Squibb Company, Foundation for the National Institutes of Health, GlaxoSmithKline, Janssen Research and Development, LLC, Lupus Foundation of America, Lupus Research Alliance, Merck & Co., Inc. Sharp & Dohme Corp., National Institute of Allergy and Infectious Diseases, National Institute of Arthritis and Musculoskeletal and Skin Diseases, Pfizer Inc., Rheumatology Research Foundation, Sanofi and Takeda Pharmaceuticals International, Inc.) created to develop new ways of identifying and validating promising biological targets for diagnostics and drug development Funding was provided through grants from the National Institutes of Health (UH2-AR067676, UH2-AR067677, UH2-AR067679, UH2-AR067681, UH2-AR067685, UH2-AR067688, UH2-AR067689, UH2-AR067690, UH2-AR067691, UH2-AR067694, and UM2-AR067678).

## Notes

### Competing Interest Statement

The authors have declared no competing interest.

